# Mechanism of N-Myc Downstream Regulated Gene-1 regulating Integrin expression to inhibit gastric cancer metastasis

**DOI:** 10.1101/2024.10.23.619963

**Authors:** Tengkai Wanga, Wenlong Ma, Di Zhang, Fenxiao Lv, Yunqing Zeng, Yuan Zheng, Mingru Liu, Jiaoyang Lu

**Affiliations:** Department of Gastroenterology, Qilu Hospital, Cheeloo College of Medicine, Shandong University, Jinan, China; Medical Integration and Practice Center, Cheeloo College of Medicine, Shandong University, Jinan, China; Department of Medical Oncology, Qilu Hospital of Shandong University, Jinan, Shandong, China

**Keywords:** Gastric cancer (GC), NDRG1, Fibronectin (FN1), Integrin

## Abstract

**Background:** N-myc downstream regulated gene-1 (NDRG1), the first member of the NDRG family, is widely distributed in various tissues and organs. In gastric cancer, the hypo-expression of NDRG1 has been positively associated with malignant biological behavior and poor prognosis. In-depth investigation of the potential regulatory mechanism of NDRG1 on gastric cancer is crucial for understanding the pathogenesis of this disease, identifying new therapeutic targets, and developing personalized treatment strategies. This study aims to explore the significance of NDRG1 in regulating the expression of integrins during the progression of gastric cancer.

**Methods:** The expression of NDRG1 in gastric cancer was analyzed using R software v4.0.3, Boxplot, and the Kaplan-Meier Plotter. Stable cell lines with NDRG1 knockdown and overexpression were subjected to wound healing assays, transwell migration assays, CCK8 assays, and colony formation assays, in order to determine the effects of NDRG1 on the migration and proliferation of gastric cancer cells. Western blotting was employed to examine the impact of NDRG1 on the expression of integrin-related proteins. Immunostaining was used to comprehensively analyze the in vitro experimental results.

**Results:** The mRNA and protein levels of NDRG1 were downregulated substantially in gastric cancer, which was inversely correlated with the malignancy of pathological behavior and poor prognosis in gastric cancer. Knockdown of the NDRG1 gene significantly enhanced the proliferation and migration capabilities of gastric cancer cells, whereas overexpression of NDRG1 reduced their metastatic potential and inhibited both migration and proliferation. NDRG1 was found to regulate the expression of fibronectin 1, which in turn influenced the expression of a series of integrins, including αv, α5, β1, and β3. This regulatory mechanism may play a crucial role in the early distant metastasis observed in gastric cancer.

**Conclusions:** This study demonstrated that NDRG1 could regulate the expression levels of integrins through specific signaling pathways, thereby interfering the adhesive and migratory capacities of gastric cancer cells. NDRG1 could serve as a valuable biomarker for the diagnosis and prognosis of gastric cancer, offering new potential targets for its treatment.

## Introduction

Gastric cancer remains one of the most prevalent malignancies globally, representing a significant threat to public health. As reported in 2022, approximately 968,000 new cases of gastric cancer and 660,000 associated deaths were recorded worldwide, underscoring its prominent position among cancers in terms of both incidence and mortality^1^. The burden of gastric cancer is particularly pronounced in China, where it ranks as the fifth most common cancer in terms of incidence and the third leading cause of cancer-related mortality, with hundreds of thousands of new cases and deaths annually^2^. The heterogeneity of gastric cancer is notable, with a wide spectrum of clinical manifestations. Its often asymptomatic nature in the early stages leads to late-stage diagnoses in a substantial number of patients, resulting in poor prognostic outcomes^3^. The pathogenesis of gastric cancer is complex and multifactorial, involving a combination of environmental exposures, dietary patterns, genetic susceptibilities, and other factors. Advances in molecular biology and genomics have facilitated the identification and clinical application of numerous gastric cancer-associated molecular biomarkers, thereby enhancing the precision of diagnostic and therapeutic approaches^4^. Notable biomarkers, such as human epidermal growth factor receptor 2 and programmed death-ligand 1^5^ have been integrated into targeted and immunotherapeutic strategies against gastric cancer^6^. The emergence of antibody-drug conjugates has further revolutionized the landscape of precision oncology in gastric cancer, contributing to a reduction in systemic toxicity while enhancing therapeutic specificity^5,7^. Thus, the ongoing discovery and validation of novel therapeutic targets for gastric cancer remain imperative, providing a robust foundation for the development of innovative targeted therapies and immunotherapeutic agents. N-myc downstream regulated gene-1 (NDRG1), the first identified member of the NDRG family, is closely linked to cellular responses to hypoxia and stress^8^. NDRG1 is ubiquitously expressed across various tissues and organs, playing a significant role in processes such as stress response, hormone regulation, cellular growth, and differentiation^9^. Emerging evidence indicates that the expression of NDRG1 varies across different tumor, where it has been implicated in the suppression of tumorigenesis and progression in cancers such as lung, breast, and colorectal cancer^10^. NDRG1 exerts its tumor-suppressive effects primarily through the inhibition of multiple oncogenic signaling pathways^11^. In the context of gastric cancer, low NDRG1 expression has been strongly associated with aggressive tumor behavior and poor clinical outcomes, implying that alterations in its expression may be intricately linked to the initiation, progression, metastasis, and prognosis of the disease^12^. Thus, elucidating the molecular mechanisms underlying NDRG1 regulation and its function in gastric cancer is of paramount importance for advancing our understanding of gastric cancer pathogenesis, discovering novel therapeutic targets, and devising individualized treatment strategies. Integrins are also known as integrins and serve as essential adhesion receptors on the surface of vertebrate cells^13^, relying on Ca2+ or Mg2+ to mediate heterotypic adhesion between cells. Integrins not only facilitate interactions between cells and the extracellular matrix (ECM) but also play a critical role in cell migration and the maintenance of tissue homeostasis^14^. In tumors, aberrant activation of integrins can promote tumor formation, growth, and metastasis^15^. High integrin expression has been observed in various cancer types, with numerous functions in tumor initiation and progression already established^16^, though there is little literature directly linking integrins to gastric cancer. Therefore, this study explores whether NDRG1 has any influence on integrins.

In this study, we identified the roles of NDRG1 in gastric cancer migration, proliferation, and colony formation. The correlation between NDRG1, fibronectin, and integrins was also confirmed. Understanding the role of NDRG1 in gastric cancer may provide deeper insights into its pathogenesis and offer potential therapeutic targets for future treatment strategies.

## 2. Materials and methods

### 2.1 Cell culture

Four human gastric cancer cell lines, including HGC-27, AGS, MKN-45, and MGC-803, as well as one immortalized normal gastric cell line, GES1, were used in this study.

Among them, HGC-27 and GES1 (Shanghai Fuheng Biotechnology) were cultured at 37°C in an environment containing 5% carbon dioxide (CO2) using DMEM medium, which was supplemented with 10% fetal bovine serum (FBS) and 1% penicillin-streptomycin. AGS (Qilu Hospital of Shandong University (Qingdao)) was cultured in a 37°C, 5% CO2 incubator using Ham’s F-12K complete medium supplemented with 10% fetal bovine serum (Gibco, AUS) and 1% penicillin-streptomycin. MKN-45 and MGC-803 (Wuhan Promoter Biotechnology) were cultured at 37°C in a 5% CO2 environment using 1640 medium, with the medium containing 10% fetal bovine serum and 1% penicillin-streptomycin.

### 2.2. Stable transfection using lentiviral infection

Use NDRG1 over express lentiviral (Ubi-MC3-3FLAG-CBh-gcGFP-IRES-puromycin) and NDRG1 reduce expression lentiviral (hU6-MCS-CBh-gcGFP-IRES-puromycin). The over express lentiviral were transfected into HGC-27 cells with HiTransG A, and the reduce expression lentiviral were transfected into MKN-45 cells with HiTransG P.The transfected cells were selected by puromycin for at least 5 days. To obtain stable control cell lines, gastric cancer cells were transfected with empty lentiviral vectors.

### 2.3 RNA extraction and real-time PCR (qPCR)

Total RNA was extracted from cultured cells with Trizol reagent, as described by the manufacturer (Invitrogen, Carlsbad, CA, USA). cDNA was synthesized from total RNA by using the Expand Reverse Transcriptase Kit (Yeasen, China). Real-time PCR was performed using cDNA in a volume of 20 μl containing 10 μl SYBR Green (Vazyme, China), 1 μl of each primer, 8.2 μl of DEPC water. PCR amplification condition was as follows: 95 °C for 30 s, 40 cycles of 95 °C for 5 s, 60 °C for 30 s. Primer for NDRG1 as 5′-GTCTCCTCTGACTTCAACAGCG -3′ (sense), 5′-ACCACCCTGTTGCTGTAGCCAA -3′ (antisense), Primer for Fibronectin as 5′-AGCCGAGGTTTTAACTGCGA -3′ (sense), 5′-CCCACTCGGTAAGTGTTCCC -3′ (antisense),Primer for β1 as 5′-GGATTCTCCAGAAGGTC -3′ (sense), 5′-TGCCACCAAGTTTCCCATCTCC -3′ (antisense), Primer for β3 as 5′-CATGGATTCCAGCAATGTCCTCC -3′ (sense), 5′-TTGAGGCAGGTGGCATTGAAGG -3′ (antisense), Primer for αV as 5′-AGGAGAAGGTGCCTACGAAGCT -3′ (sense), 5′-GCACAGGAAAGTCTTGCTAAGGC - 3′ (antisense), Primer for α5 as 5′- CGGGCTCCTTCTTCGGATT -3′ (sense), 5′- CACCCCAAGGACAGAGGTAG -3′ (antisense),GADPH as 5′- GTCTCCTCTGACTTCAACAGCG-3′ (sense), 5′-ACCACCCTGTTGCTGTAGCCAA-3′ (antisense). As negative control, ddwater was run with every PCR.

### 2.4. Western blotting analysis

Protein was extracted using SDS lysis buffer and transferred to PVDF membranes. After the membranes were blocked with 5% skim milk for 2 hours, they were incubated with primary antibodies overnight at 4 °C, which was followed by incubation with secondary antibodies for 1 h. Bands were visualized using GE healthcare (AI 680) and analyzed with ImageJ software. GAPDH (ET1601-4, 1:10000; Huabio) was used as a protein loading control. Primary antibody against NDRG1 (ab124689, 1:5000; abcam), Fibronectin (ab2413, 1:1000; abcam),alphaV (ab179475, 1:5000; abcam),alpha5 (ab150361, 1:5000; abcam),Beta 1 (AB179471, 1:1000; abcam), Beta 3 (AB179473, 1:1000; abcam).All secondary antibodies were purchased from Zhongshan Golden Bridge Biotechnology Company Limited.

### 2.5. Wound-healing assay

Cells were seeded in 6-well plates. When cells reached confluency, a wound was created using a 1000-μL sterile pipette tip and photographed (0 h). The rate of gap closure was measured at next day same time (24 h). Each experiment was performed three times.

### 2.6. Cell invasion assay and cell migration assays

For migration assays, HGC-27 cells (1 × 105) were suspended in serum-free medium and added 200μL to the upper chamber of the Transwell plate (Corning Incorporated, USA). DMEM with 10% FBS was added to the bottom chamber in 24-well plates. After the cells were incubated for 24 h, they were fixed with methanol and stained with crystal violet for 15 min.MKN-45 cells (1 × 105) were suspended in serum-free medium and added to the upper chamber of the Transwell plate(Corning Incorporated, USA). 1640 with 10% FBS was added to the bottom chamber in 24-well plates. After the cells were incubated for 72 h, they were fixed with methanol and stained with crystal violet for 15 min. Invasion assays were performed the same say as the migration assays except that the Transwell chambers were coated with Matrigel before the cells were seeded in the upper chamber. These cells were counted using an inverted light microscope (Nikon). Each experiment was performed three times.

### 2.7 Cell proliferation and colony formation assay

HGC-27 and MKN-45 cells were inoculated into 96-well plates at a density of 2500 cells per well. Cell proliferation was measured at specific time points using the Cell Counting Kit-8 (Beyotime). Following the provided, 100 µL of serum-free medium with 10% CCK-8 reagent was added to each well and the plates were incubated at 37 °C for 2 hours. The absorbance values were measured at 450 nm using a microplate reader(), and growth curves were plotted based on absorbance and time. For the cell colony formation assay, 1000 HGC-27 cells and 1000 MKN45 cells were inoculated in 6-well plates and cultured for 10—14 days. Afterward, the cells were fixed using 4% paraformaldehyde and stained with 0.1% crystal violet. Observe the cloning rate and clone size.

### 2.8. Immunohistochemistry (IHC)

Immunohistochemistry (IHC) was performed according to the manufacturer’s constructions. After the samples were deparaffinized with xylene and rehydrated with ethanol, the samples were incubated with 3% H2O2 for 5 min to block endogenous peroxidase activity. Then, boiled in 0.01 M citrate buffer (pH 6.0) for 20 min at 95 °C for antigen recovery. Then blocked the samples with 5% normal goat serum for 20 min at 20 °C. Subsequently, the sections were incubated with polyclonal antibodies against NDRG1(1:500, ab124689, Abcam) at 4 °C overnight and then incubated with secondary antibodies(1:200, Servicebio, Wuhan, China).The results were imaged under a fluorescence microscope (Olympus, Japan).

### 2.9. Immunofluorescence (IF)

HGC-27, MKN-45 were seeded on coverslips 48 h. After treatment, cells were fixed with 4% paraformaldehyde for 15 min. Fixed and permeabilized cells were then washed with PBS thrice and incubated with 2% Normal Horse Serum (NHS) for 30 min. Then incubated with primary antibodies overnight at 4 °C, After further washes, secondary anti-rabbit antibody tagged to Alexafluor 594 (1:1000 dilution in NHS) along with Phalloidin tagged to Alexafluor 488 (1:500 dilution in NHS) and DAPI was added to the coverslips and incubated for 10 min. Coverslips were then washed and mounted on glass slides using mounting medium. Cells were visualized under inverted fluorescence microscope (Olympus, Japan) under blue, red and green channels. Images were quantified by measuring the ratio of fluorescence in the nuclear region to that in the total cell using ImageJ software.

### 2.10. Statistical analysis

All data were expressed as mean ± SD. Statistical analyses were performed using Prism software (GraphPad Software 8), and consisted of analysis of variance followed by Student’s t-test when comparing two experimental groups. One-way ANOVA analysis was used to compare the differences among groups. Overall survival (OS) rates were determined using the Kaplan-Meier method. All experiments were triplicated, and p < 0.05 was considered significant.

## 3. Results

### 3.1. Expression of NDRG1 in gastric cancer tissues and cells with survival analysis

We initially retrieved RNA sequencing data for gastric cancer and the corresponding clinical information from The Cancer Genome Atlas (TCGA) dataset. The expression of NDRG1 was significantly lower in gastric cancer tissues compared to normal controls (Fig. 1A). Similar results were corroborated using datasets from the GEO database (Fig. 1B). Further validation through immunohistochemistry and immunofluorescence staining of normal and cancerous gastric tissues confirmed reduced NDRG1 expression in gastric cancer tissues relative to normal tissues (Fig. 3C, D). Leveraging clinical data from the TCGA, we generated scatter plots, gene expression heatmaps, and survival curves, revealing that low NDRG1 expression was significantly associated with poorer prognosis in gastric cancer. The median survival time in the low NDRG1 expression group was 2.2 years, compared to 3.4 years in the high expression group. Ranking samples by NDRG1 expression levels demonstrated that lower NDRG1 expression corresponded with higher mortality rates and shorter survival durations (Fig. 1E-G). Additionally, we assessed NDRG1 mRNA and protein levels in three widely studied gastric cancer cell lines (Fig. 1H, I).To further investigate the diagnostic value of NDRG1 in GC, we applied the TCGA dataset to plot NDRG1 time-dependent ROC curves to predict survival in GC patients at 1, 3 and 5 years. The results showed that 3 and 5 years AUC values were >0.5 (Fig. 1G), shows that it is very meaningful to predict the survival rate of GC. These results suggested that NDRG1 has the potential to become a diagnostic marker for GC.

### 3.2. Overexpression of NDRG1 inhibit the invasion, migration and proliferation of GC cell lines

Based on the experimental results from the previous phase, the HGC27 cell line exhibited the lowest expression level of NDRG1. Therefore, we selected the HGC27 gastric cancer cell to construct an NDRG1 overexpression cell strain for further investigation. The overexpression efficiency is shown in the figure(Fig.1A,B).The cell proliferation of HGC27-OE in the CCK8 assay was significantly slower than that in the control group(Fig.2C). The Wound-healing assay show that the healing speed of the HGC27 OE was significantly slower than in the control group(Fig.2D).Compared with the control group, the colony formation rate and colony formation size of HGC27 cells in the HGC27 OE group in the cloning experiment were significantly reduced(Fig.2E). The number of HGC27 OE cells in the Transwell experiment was significantly less than that in the control group(Fig.2F).

**Fig. 1.**
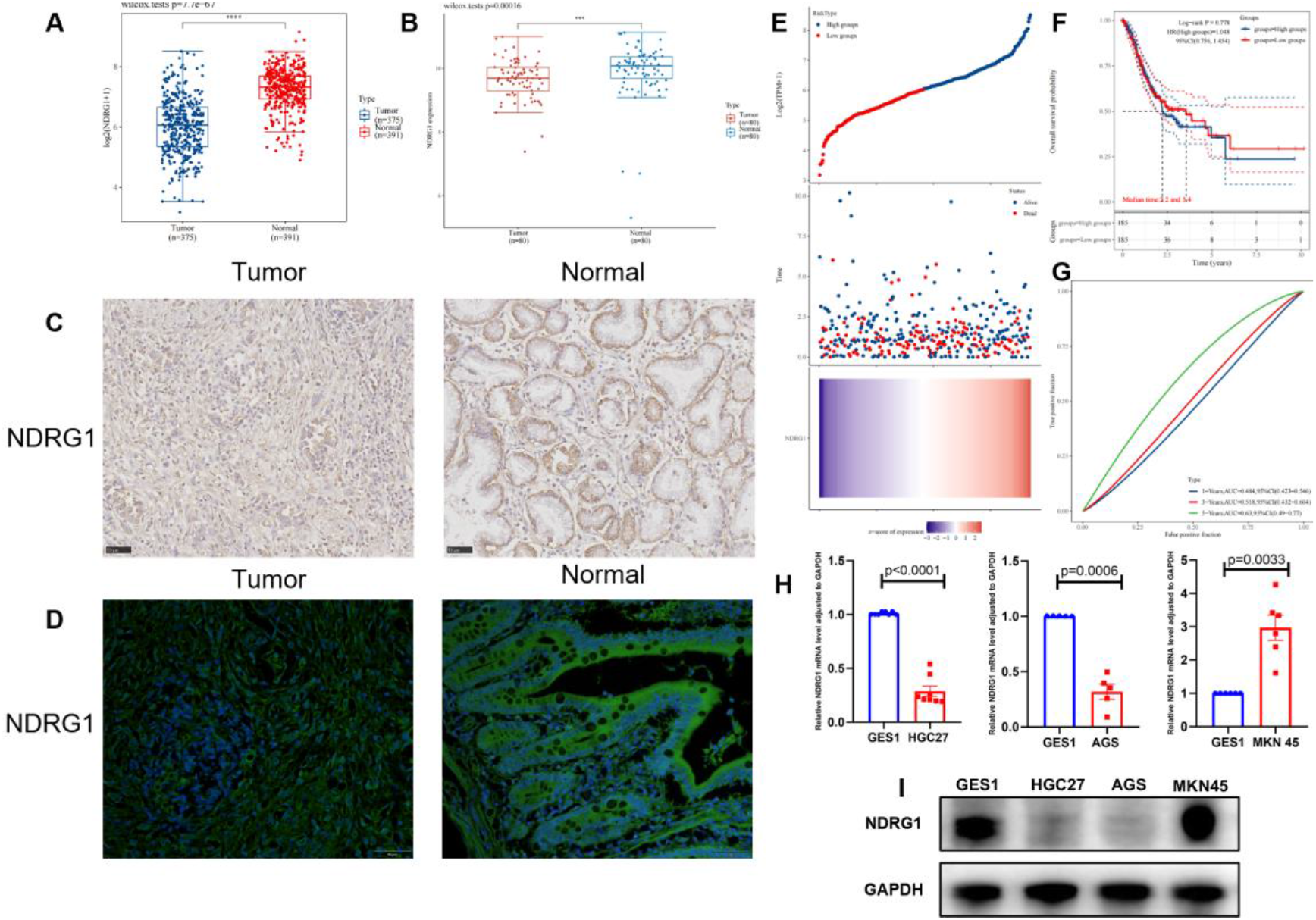
NDRG1 expression in GC tissues and cell lines. (A). The NDRG1 expression in TCGA database. (B). The NDRG1 expression in GEO database. (C). The NDRG1 expression in GC and adjacent normal tissue use Immunohistochemistry assay. (D). The NDRG1 expression in GC and adjacent normal tissue use Immunofluorescence assay. (E) scatter plot of NDRG1 expression and patient survival; (F) correlation between NDRG1 expression and the prognosis of GC; (G) the diagnostic value of NDRG1 in GC based on TCGA data; (H, I)expression of NDRG1 in three gastric cancer cell lines and normal gastric mucosal epithelial cell.(All data are presented as mean ± SEM. A t-test was used for statistical analysis. the p value < 0.05 was considered to indicate statistical significance.)

**Fig. 2.**
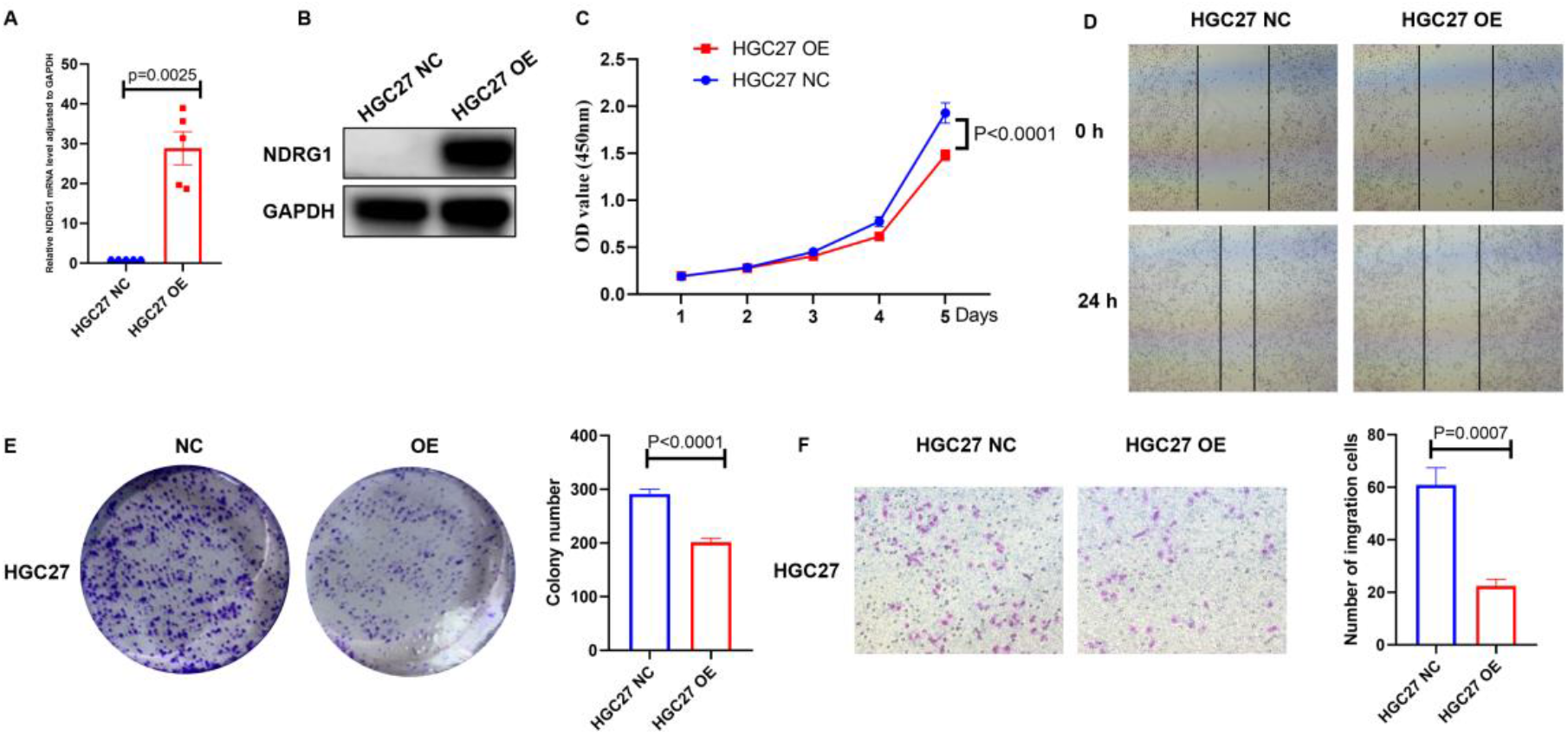
Overexpression of NDRG1 inhibit the invasion, migration and proliferation of GC cell lines. (A,B) Expression of NDRG1 in HGC27 OE stably transfected GC cell line; (C) the proliferation curve of HGC27 NC and HGC27 OE cells; (D) wound healing results for HGC27 NC and HGC27 OE cells; (E) results of colony formation experiments for HGC27.(F) HGC27 NC and HGC27 OE transwell results and statistical chart of the number of migration experiments. (All data are presented as mean ± SEM. A t-test was used for statistical analysis. the p value <0.05 was considered to indicate statistical significance.)

### 3.3. Knockdown of NDRG1 promotes invasion, migration, and proliferation in GC cell line

The MKN45 cell line exhibited high NDRG1 expression, so we selected MKN45 cell line to further study the role of NDRG1 in gastric cancer cells. NDRG1 reduce expression lentiviral were transfected into MKN45 cells with HiTransG P. After the transfected cells were selected by puromycin for at least 5 days. The knockdown efficiency is shown in the figure(Fig. 3A,B). The cell proliferation of MKN45 96039 in the CCK8 assay was significantly faster than that in the control group(Fig. 3C). The Wound-healing assay show that the healing speed of the MKN45 96039 was significantly faster than in the control group(Fig. 3D). The colony formation rate and colony formation size of MKN45 cells in the NDRG1-Si group in the cloning experiment were significantly higher than those in the control group(Fig. 3E).The number of MKN45 96039 cells in the Transwell experiment was significantly greater than that in the control group. (Fig. 3F).

**Fig. 3.**
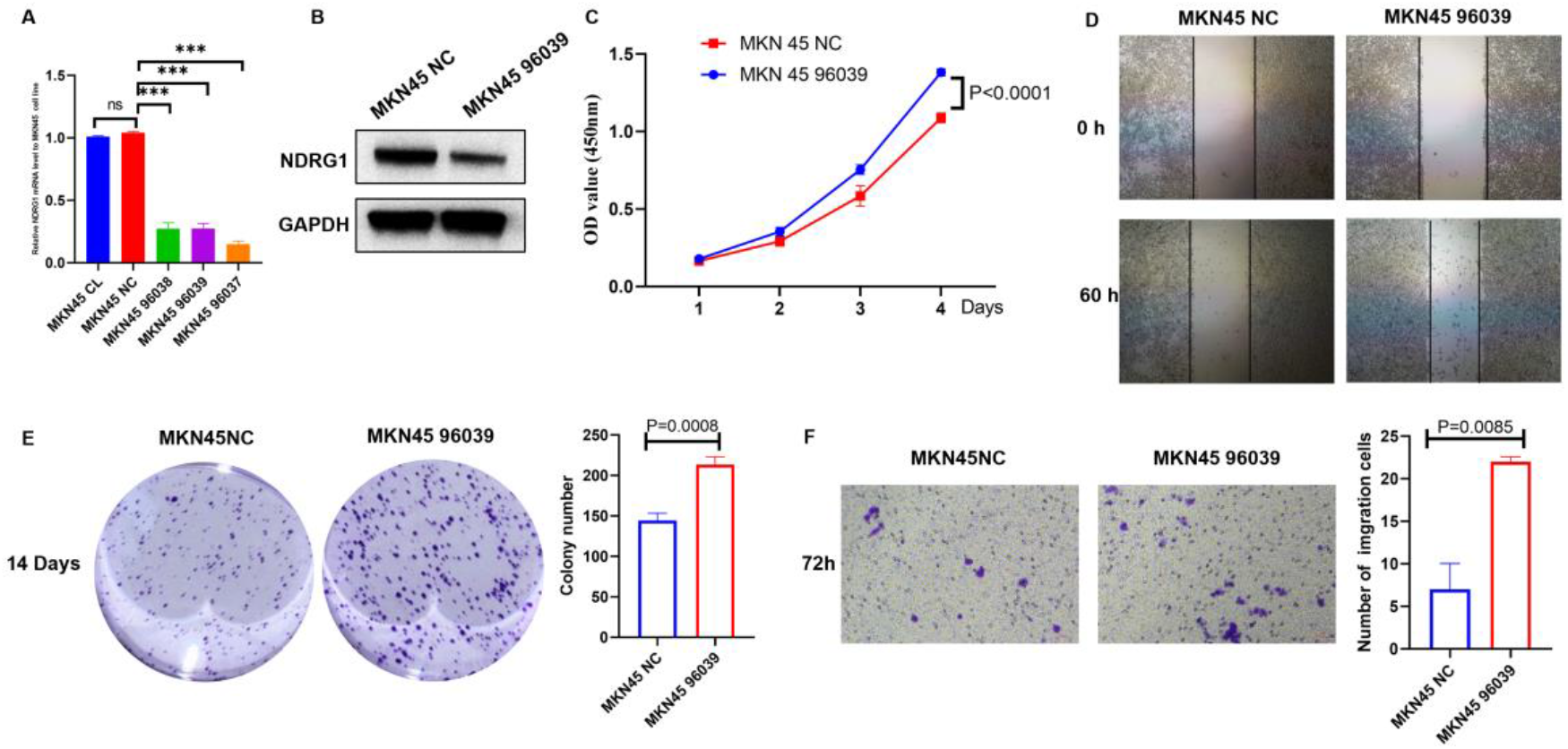
Knockdown of NDRG1 promotes the invasion, migration and proliferation of GC cell lines. (A,B) expression of NDRG1 in MKN45 NC and MKN45 96039 stably transfected gastric cancer cell line; (C) the proliferation curve of MKN45 cells; (D) wound healing results for MKN45 cells; (E) results of colony formation experiments for MKN45. (H) MKN45 transwell results and statistical chart of the number of migration experiments; (All data are presented as mean ± SEM. A t-test was used for statistical analysis. the p value <0.05 was considered to indicate statistical significance.)

### 3.4. Knockdown of NDRG1 promotes adhere and the adherent area increases in GC cells

In a fortuitous observation that we found some morphological differences between NDRG1 Overexpression cells and NDRG1 knockdown cells. HGC27 and MKN45 cells were plated on plates with tissue culture treated and allowed to spread, cells were visualized by microscope for their ability to spread. After 12 hours, as seen in the photomicrograph, MKN45 96039 cells stick to the plates faster than MKN45 NC cells. MKN45 NC cells remained rounded and had difficulties to spread(Fig. 4A). The opposite result was observed for HGC27 cells in the NDRG1-OE group(Fig. 4B). These data suggest that NDRG1 knockdown cells exhibit increased kinetics of cell spreading.

**Fig. 4.**
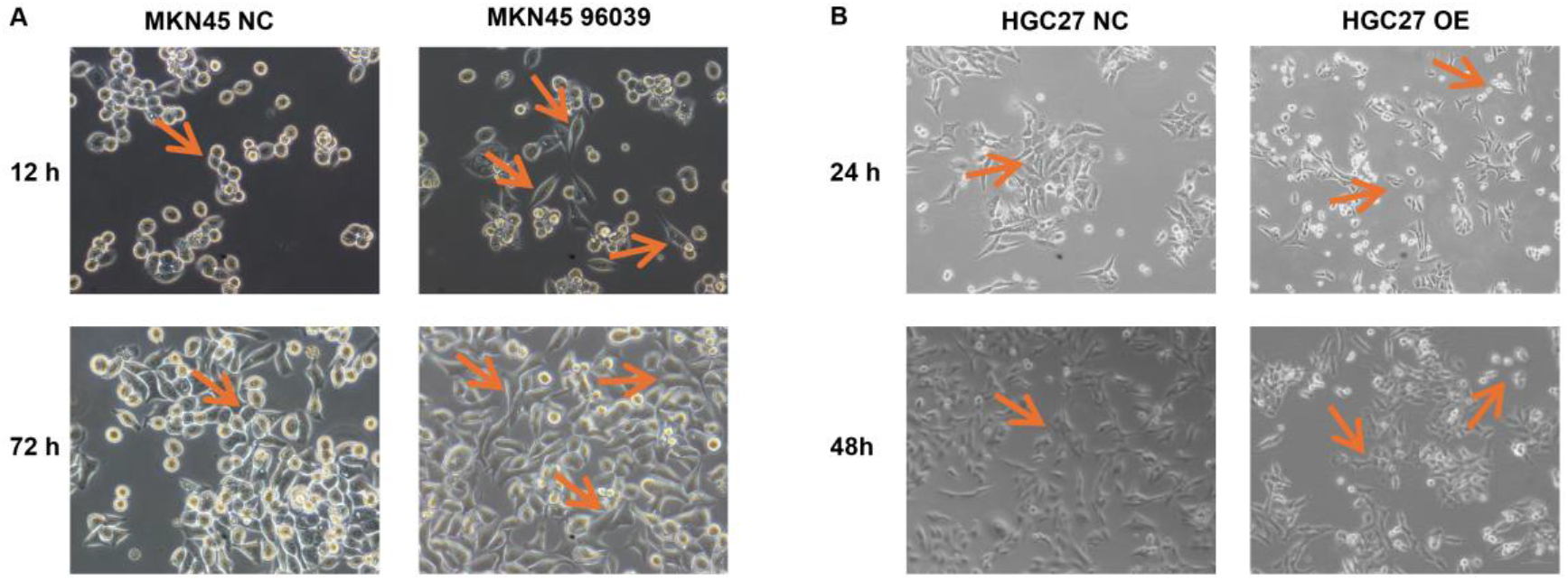
Knockdown of NDRG1 promotes adhere and the adherent area increases in GC cells.(A) MKN45 96039 cells began to exhibit extensive adherence to the substratum at 12-hour, within the same timeframe, most of the MKN45 NC cells still remained rounded. After 72 hours of cell culture, the majority of MKN45 96039 cells exhibit a spindle shaped morphology. (B) After 24 hours of cell culture, the adherent area of HGC27 OE cells remains relatively small. After culturing HGC27 OE cells for 48 hours, some HGC27 OE cells still maintained a round shape.(All experiments were repeated at least three times.)

### 3.5. Overexpression of NDRG1 increased the expression of fibronectin1 and integrins in GC cells

Cell spreading occurs due to coordinated action of integrin-mediated adhesion to ECM and subsequent remodeling of the actin cytoskeleton. To investigate whether knockdown of NDRG1 affects cellular adhesion, we performed qPCR assay and Immunofluorescence assay on HGC27 NC and HGC27 OE cells. We investigated the regulatory role of fibronectin1 in NDRG1 gene transcription. The Overexpression of NDRG1 significantly increased fibronectin1 expression at both the mRNA and protein levels (Fig. 5A,B). Immunofluorescence experiments revealed a significant increase in FN1 protein levels after 48 h of cell cultured(Fig. 5C). That is not all, Overexpression of NDRG1 can also increase Integrin α5, αv, β1, β3 expression at mRNA level (Fig. 5D). Immunofluorescence also experiments revealed a significant increase in α5, αv, β1, β3 protein levels after 48 h of cell cultured(Fig. 5E).

**Fig. 5.**
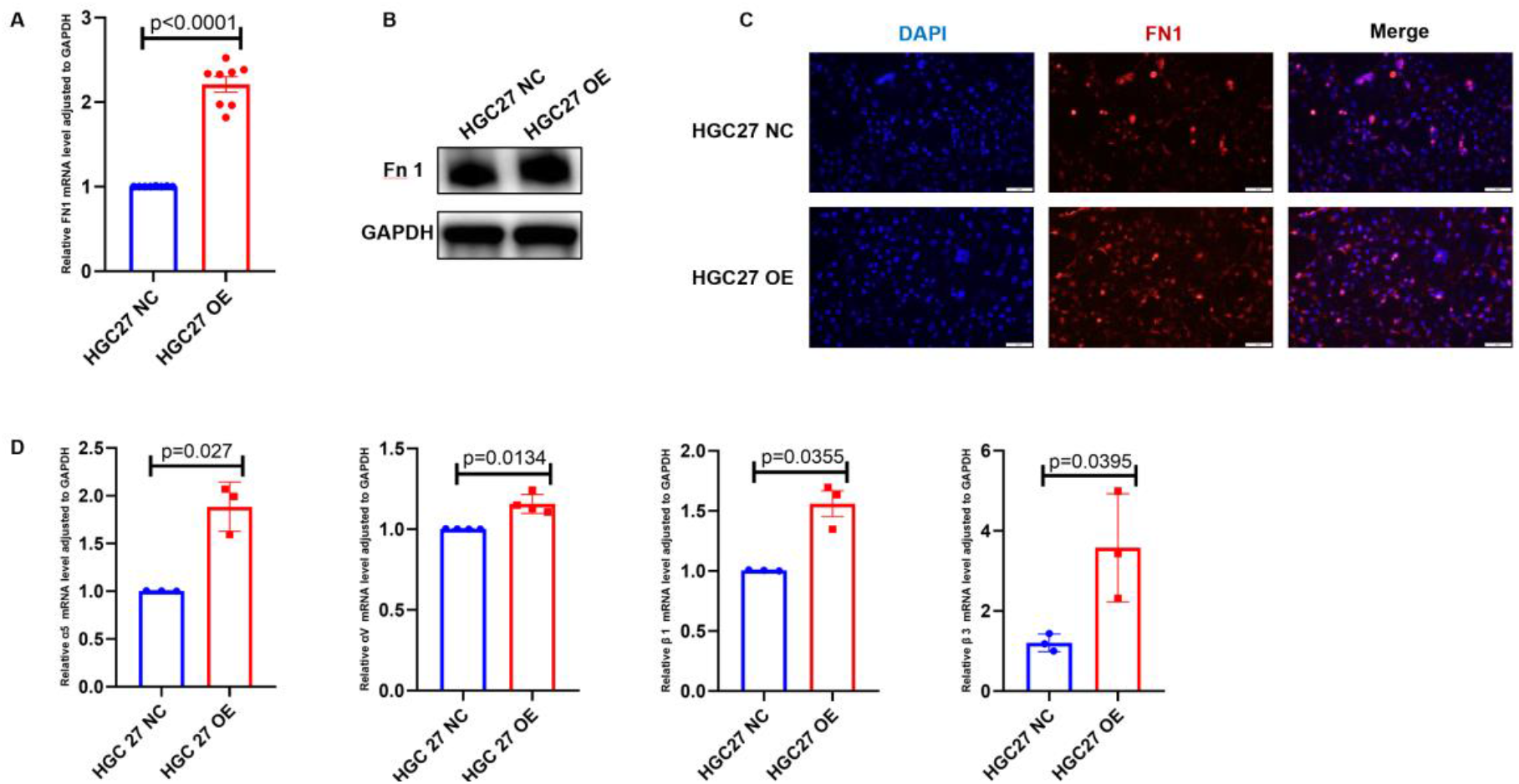
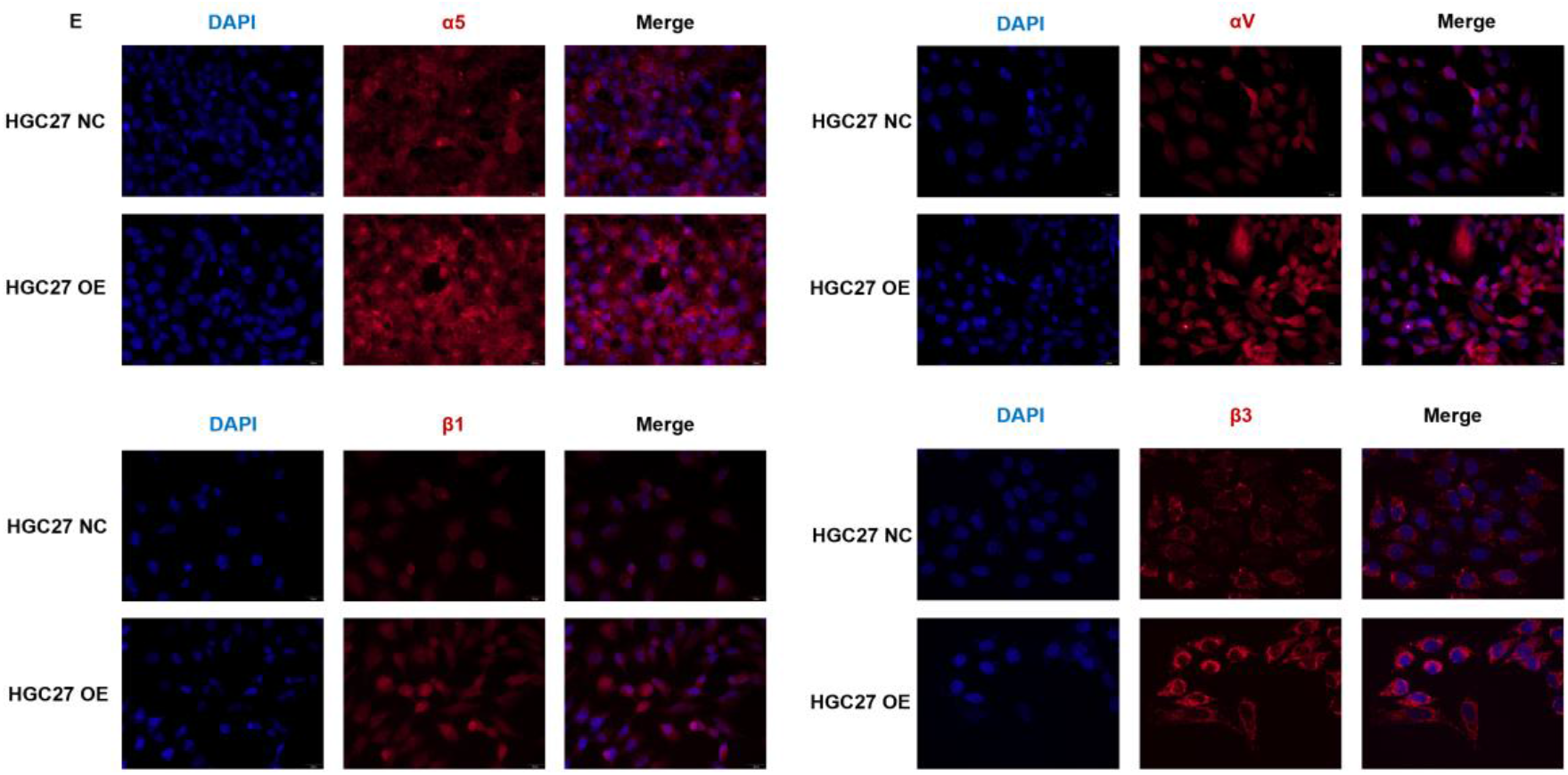
NDRG1 represses the transcription of fibronectin1 and integrins in HGC27 OE cells. (A)mRNA level of FN1 gene in NDRG1 Overexpression cells. (B) Immunoblotting analysis of FN1 protein levels in NDRG1overexpression cells. (C) Immunofluorescence analysis of FN1 protein in HGC27 NC and HGC27 OE cells cultured in plates for the indicated time. (D)mRNA level of α5, αv, β1, β3 in NDRG1 Overexpression cells. (E) Immunofluorescence analysis of α5, αv, β1, β3 protein in HGC27 NC and HGC27 OE cells cultured in plates for the indicated time. (All data are presented as mean ± SEM. A t-test was used for statistical analysis. the p value <0.05 was considered to indicate statistical significance.)

### 3.6. Knockdown of NDRG1 represses the expression of fibronectin1 and integrins in GC cells

We selected MKN45 GC cell for further investigation due to its significant alteration upon NDRG1 knockdown. We investigated the regulatory role of NDRG1 in FN1, α5, αv, β1, β3 gene transcription. The knockdown of NDRG1 significantly decreased FN1, α5, αv, β1, β3 expression at both the mRNA and protein levels (Fig. 6A,B,D,E). Immunofluorescence experiments also revealed a significant decrease in FN1 protein levels after 48 h of cell cultured (Fig. 6C).

**Fig. 6.**
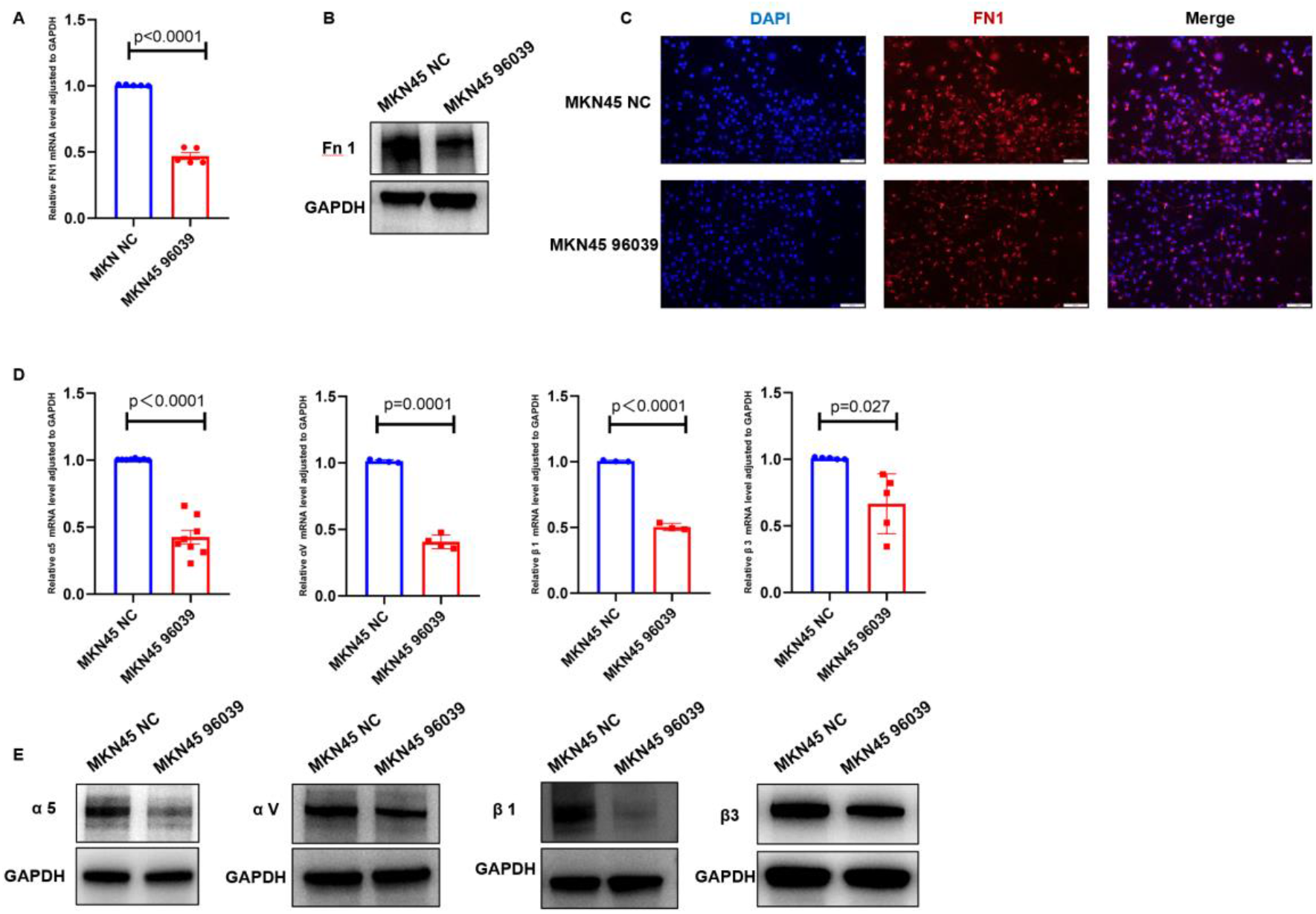
Knockdown of NDRG1 represses the transcription of fibronectin1 and integrins in MKN45 cells. (A)mRNA level of FN1 gene in NDRG1 Knockdown cells. (B) Immunoblotting analysis of FN1 protein level in NDRG1 Knockdown cells. (C) Immunofluorescence analysis of FN1 protein in MKN45 NC and MKN45 96039 cells cultured in plates for the indicated time. (D)mRNA level of α5, αv, β1, β3 in NDRG1 Knockdown cells. (E)Immunoblotting analysis of α5, αv, β1, β3 protein levels in NDRG1 Knockdown cells. (All data are presented as mean ± SEM. A t-test was used for statistical analysis. the p value <0.05 was considered to indicate statistical significance.)

## Discussion

NDRG1 has been extensively investigated as a prognostic marker and predictor of tumor progression across various cancers. In hepatocellular and lung carcinomas, elevated expression of NDRG1 has been linked to poor clinical outcomes, with evidence suggesting its involvement in tumor angiogenesis and unfavorable prognoses in these malignancies. Within the context of breast cancer, NDRG1 expression is notably higher in the estrogen receptor-negative subtype, serving as an independent prognostic factor for adverse survival in patients with inflammatory breast cancer^17-19^. Conversely, a substantial body of research indicates that NDRG1 functions as a metastasis suppressor in breast cancer; its expression is significantly diminished in metastatic cases, particularly in those with lymphatic or bone dissemination, compared to patients with localized disease^19-21^. In oral squamous cell carcinoma, NDRG1 deficiency is associated with regional metastasis, primarily through the induction of epithelial-mesenchymal transition^22^. Despite the heterogeneous roles of NDRG1 across different tumor types, it remains a promising therapeutic target and biomarker for predicting cancer prognosis. As for gastric cancer, prior studies demonstrated the tumor-suppressive effects of NDRG1, specifically in inhibiting tumor invasion and migration. Furthermore, hypo-expression of NDRG1 was reported correlating with increased malignancy and poorer prognosis in gastric cancer^12,23^, which was consistent with our bioinformatics analysis results. NDRG1 exhibits multifaceted roles in tumor biology, particularly as a well-recognized metastasis suppressor. Tumor cell migration is a critical step in the metastatic cascade and is frequently accompanied by alterations in the ECM during malignant progression^24^. NDRG1 has been shown to modulate tumor cell migration by influencing the interactions between cells and the ECM. Fibronectin, a key ECM component, has a dual role in tumor biology that is often context-dependent. While matrix-associated fibronectin is generally regarded as a facilitator of tumor metastasis, numerous studies have highlighted its tumor-suppressive functions, particularly when derived from tumor cells themselves^25,26^. In many metastatic cancer cells, fibronectin expression is either significantly reduced or entirely absent^27^. Reinstating fibronectin expression in metastatic rat renal cells has been shown to restore normal cell monolayer morphology^28^, while overexpression of recombinant fibronectin effectively suppresses the metastatic phenotype in human fibrosarcoma cells^29^. Our observations reveal that gastric cancer cell lines with upregulated NDRG1 exhibit decreased migratory capacity alongside elevated fibronectin expression, suggesting a potential mechanism through which NDRG1 modulates gastric cancer cell migration via fibronectin.

Fibronectin typically binds to cells via integrins, thereby influencing cell migration ability. Integrins are heterodimers formed by α and β subunits, with integrins α5, αv, β1, and β3 being most closely associated with fibronectin. In Chinese hamster ovary cells, overexpression of the fibronectin receptor integrin α5β1 induces greater fibronectin deposition, significantly reducing the migration of these cells^30^. Loss of the α2β1 integrin increases breast cancer metastasis, while α2β1 integrin expression inhibits migration, intravasation, and anchorage-independent growth in vitro^31^. Our results indicated that, following NDRG1 overexpression, the expression of integrins α5, αv, β1, and β3 was significantly upregulated, suggesting that these integrins may inhibit gastric cancer cell migration by regulating the binding between fibronectin and cells.

In this study, we obtained RNA sequencing data and corresponding clinical information for gastric cancer from the TCGA database. Through scatter plots, gene expression heatmaps, and survival curves, we observed that low expression of NDRG1 was negatively correlated with malignant biological behavior and positively correlated with poor prognosis in gastric cancer, providing further support for the use of NDRG1 in clinical cancer prognosis. Moreover, we found that overexpression of NDRG1 could suppress tumor cell invasion and migration, while its downregulation had the opposite effect. Overexpression of NDRG1 led to higher expression of fibronectin and its related integrin receptors, suggesting that fibronectin and integrins may play a significant role in the regulatory effects of NDRG1 on tumors.

## Acknowledgments

We acknowledge the Natural Science Foundation of Shandong Province (ZR2020QH184) and the ACTA, GEO database for providing open access to their data.

## Authorship contribution statement

**Tengkai Wang:** Writing - original draft, Visualization, Validation, Methodology, Investigation, Formal analysis, Data curation, **Wenlong Ma** Writing-review & editing, Visualization, Project administration. **Di Zhang** Writing-review & editing, Resources, Project administration. **Fenxiao Lv:** Writing-review & editing, Validation. **Yunqing Zeng:** Project administration. **Yuan Zheng:** Validation. **Mingru Liu:** Data curation. **Jiaoyang Lu:** Supervision, Project administration, Funding acquisition, Conceptualization.

